# Limited roles of Piezo mechanosensing channels in articular cartilage development and osteoarthritis progression

**DOI:** 10.1101/2022.10.07.511314

**Authors:** Cameron Young, Tatsuya Kobayashi

## Abstract

Osteoarthritis (OA) is a prevalent disease characterized by degeneration of the joint and pain. Mechanical stress plays a central role in OA development. It is hypothesized that cells in the OA joints produce OA-promoting molecules upon mechanical stress, and therefore, the mechanosensing systems are a theoretical target for OA treatment. Piezo mechanosensing channels mediate high-level mechanical stress in chondrocytes and have been suggested to play an important role during OA progression. To test this hypothesis, we ablated *Piezo1* and *Piezo2* in joint tissues using *Gdf5-Cre* transgenic mice [*Piezo1* and *2* doubly conditional knockout (cKO) mice, cKO mice]. cKO mice showed normal development of knee joints. Both control and cKO mice developed modest to severe OA 12 weeks after the induction of OA, although some cKO mice showed milder OA. We did not find significant differences in pain in mice or gene expression after fluid flow stress in primary cells between control and cKO. Our data demonstrate the limited role of Piezo channels in joint development and OA progression.

**Summary:** *Objective:* To investigate the role of Piezo1 and Piezo 2 in surgically induced osteoarthritis (OA) in mice.

*Design:* Male conditional knockout (cKO) mice missing *Piezo1* and *Piezo2* in the joint via *Gdf5-Cre* transgenic mice were induced post-traumatic osteoarthritis (OA) by destabilization of the medial meniscus (DMM) of the right knee joint at 12 weeks of age. The severity of OA was assessed at 24 weeks of age using a modified Osteoarthritis Research Society International (OARSI) scoring system. OA-associated pain was evaluated by static weight bearing analysis at 4, 8, and 12 weeks post-operation. Additionally, articular chondrocytes isolated from cKO mice were exposed to fluid flow shear stress (FFSS) to evaluate the expression of OA-associated genes.

*Results:* Mice with conditional deletion of *Piezo1* and *Piezo 2* showed normal joint development with no overt histological changes in the knee joint at 12 weeks and 24 weeks. DMM surgery induced moderate to severe OA in both control and cKO mice, although a few cKO mice showed milder OA. Pain assessment by static weight-bearing analysis suggested Piezo ablation in the joint has no beneficial effects on pain. FFSS increased the expression of OA-related genes both in control and cKO mice to similar extents.

*Conclusion:* Piezo1 and Piezo2 are not essential for normal joint development. Genetic ablation of Piezo channels did not confer evident protective effects on OA progression in mice. *In vitr*o data suggests that different mechanotransducers other than Piezo channels mediate FFSS in mechanical stress-induced gene expression.

## Introduction

Osteoarthritis (OA), a common disease characterized by chronic pain and degeneration of joints, widely affects people worldwide with increasing prevalence. OA is strongly associated with aging and injury, and cumulative and/or excessive mechanical stress plays a central role in the pathogenesis of OA. In addition to direct physical damage to the cartilage, many factors, such as inflammation and cellular senescence, are known to contribute to development and progression of OA. Cells in the joint affected by OA respond to mechanical stress and secrete bioactive molecules that could further progress OA. Mechanical stress is sensed by the cell through multiple mechanisms such as primary cilium, integrins, and mechanosensing ion channels^1^. Among diverse groups of ion channels expressed in chondrocytes, two groups of mechanosensitive calcium ion channels, transient receptor potential (TRP) channels, including TRP vanilloid-4 (Trpv4), and Piezo-1 (Piezo1) and −2 (Piezo2), are particularly of interest as possible therapeutic targets for OA^2^. Trpv4 appears to regulate anabolic chondrocyte metabolism and Ca^2+^ levels in response to low strain, physiologic mechanical stress maintaining proper extracellular matrix for cartilage health ^3^, whereas the effect of its inhibition on OA is unclear. Conditional ablation of *Trpv4* reduces age-related OA-like changes, although it does not significantly alter the severity of OA induced by destabilization of the medial meniscus (DMM)^4^. Piezo channels are broadly expressed in different types of cells. Piezo1 is highly expressed in skeletal tissues, including cartilage, and is activated by high-level, injurious strain ^5^. A study using isolated chondrocytes showed that Piezo1 and Piezo2 synergistically regulate Ca^2+^ influx in response to mechanical stress ^6^. Furthermore, this study demonstrated that pharmacological Piezo inhibition reduced chondrocyte apoptosis in cartilage explants upon injurious mechanical loading. Inflammatory cytokines and matrix-degrading enzymes, such as interleukins and matrix metalloproteinases, are well-known factors that accelerate OA^7, 8^. Interleukin-1α enhances Piezo1 expression, which appears to further amplify the effect of mechanical stress to form a feed-forward loop to promote OA progression ^9^. These studies suggest that Piezo ion channels play a significant role as a mediator of mechanical stress during OA development and progression. In this study, we have tested the hypothesis that the conditional ablation of Piezo channels in the joint confers protective effects in a mouse OA model.

## Method

### Mice

Floxed *Piezo1* and floxed *Piezo2* mice were purchased from the Jackson laboratory. *Gdf5-Cre* transgenic mice were previously described^10^. Mice were in a congenic strain derived from the C57/BL6 and FBV strains. Only male mice were subjected to the study. Doubly conditional homozygous knockout males (cKO, *Gdf5-Cre:Piezo1*^*fl/fl*^*:Piezo2*^*fl/fl*^) and male Cre-negative littermate controls (*Piezo1*^*fl/fl*^*:Piezo2*^*fl/fl*^) were generated and were subjected to DMM surgery. This study was approved by the Institutional Animal Care and Use Committee (IACUC) at Massachusetts General Hospital and by the Animal Care and Use Review Office (ACURO) of the U.S. Army Medical Research and Development Command (USAMRDC).

### DMM surgery

The destabilization of the medial meniscus (DMM) surgery was performed to induce OA according to an established protocol with minor modifications ^11^. Briefly, under anesthesia, the meniscotibial ligament was cut through a 1 cm skin incision and a 0.3 cm incision of the right joint capsule. The joint capsule was then sutured with 7-0 Vicryl sutures, and the skin incision was closed using 9 mm wound clips. Wound clips were removed 14 days after the operation.

### Mouse Dissection and Histological analysis

cKO and control male mice were sacrificed at 12 weeks or 24 weeks without surgery for baseline histological assessment, and mice that received DMM surgery were sacrificed at 24 weeks (12 weeks post-operation). Right hindlimbs were fixed in 10% Formaldehyde-PBS, decalcified, paraffin-processed, sectioned, and stained with Safranin-O or hematoxylin and eosin (H/E) according to a standard procedure. Osteoarthritis (OA) was scored using the Osteoarthritis Research Society International (OARSI) mouse OA scoring system ^12^. Sagittal sections from three different cutting planes of the mid-tibial plateau per mouse were blindly evaluated by two inspectors. The average OARSI score per mouse was then calculated. Due to the difficulty of obtaining comparable sections in the femoral condyle, only the tibial plateau was evaluated.

### Articular Chondrocyte Isolation and Fluid Flow Shear Stress

Articular chondrocytes were obtained from 10-day-old mouse knee and hip joints according to the method previously described ^13^. Briefly, the tibial condyle or hip joint was manually dissected. The soft tissues and subchondral tissues were manually removed as much as possible. Cartilage pieces were digested overnight in a growth medium (DMEM containing 10% fetal bovine serum) and 0.2% collagenase II (Worthington) to disperse chondrocytes. Cells were passed through 0.40 μm nylon mesh, resuspended in the growth medium, plated in 48-well plates, and grown to confluence. Fluid flow shear stress (FFSS) was applied for 4 hours by shaking the plate at 1,200 rpm on a mixing platform (Thermomixe R, Eppendorf).

### qRT-PCR

Gene expression was analyzed by quantitative reverse transcription polymerase chain reaction (qRT-PCR). RNA was isolated from cells using the Directzol RNA Mini-Prep Kit (Zymo Research) and converted to cDNA using the Verso cDNA synthesis kit (Thermo Scientific). qRT-qPCR was performed using the PerfeCTa SYBR Green Supermix (QuantaBio). The values were normalized to *Actb*. PCR primer sequences are as follows: *Actb*-L, 5’- GCACTGTGTTGGCATAGAGG −3’ and *Actb*-R, 5’- GTTCCGATGCCCTGAGGCTCTT −3’; *Piezo1*-L, 5’- GATTGGGCAGCGTATGAACT −3’ and *Piezo1*-R, 5’-GTACAGCAGGAACAGCGTGA −3’; *Piezo2*-L, 5’- CCTCGTGTTTGGGATTCACT −3’ and *Piezo2*-R, 5’- GTAGCCACAGCGGATTTGAT −3’; *Acan*-L, 5’-GAAGAGCCTCGAATCACCTG-3’ and *Acan*-R, 5’- ATCCTGGGCACATTATGGAA −3’; *Mmp13*-L, 5’- GCCATTTCATGCTTCCTGAT −3’ and *Mmp13*-R, 5’-TTTTGGGATGCTTAGGGTTG −3’; *Wnt11*-L, 5’--CAGGATCCCAAGCCAATAAA −3’ and *Wnt11*-R, 5’- GACAGGTAGCGGGTCTTGAG −3’; *Il6-*L, 5’- CACAAGTCCGGAGAGGAGAC −3’ and *IL6*-R, 5’- TCCACGATTTCCCAGAGAAC −3’.

### Pain assessment and OA scoring

The OA-associated pain was assessed using an incapacitance meter (BioSeb) at 4, 8 and 12 weeks after the DMM surgery. The weight distribution difference between the injured (right) and unaffected (left) hind limb was quantified to evaluate pain. The average value of ten measurements for each mouse at each time point was calculated.

### Statistical analysis

Statistical analysis was performed using Graphpad Prism 9.4. For two-group comparisons, two-tailed, Mann-Whitney *U* test was used to compare OA and pain assessment. Student *t*-test was used for other analyses.

## Results

To confirm the deletion of *Piezo1* and *Piezo2*, articular chondrocytes were isolated from the knee and hip joints of cKO and control mice. Relative expression of *Piezo1* and *Piezo2* was decreased by 50 −75% in the hip and knee joint articular chondrocytes in cKO mice (Fig. 1A). This partial reduction is likely due to contamination of non-articular chondrocytes including growth plate chondrocytes that are difficult to eliminate by manual dissection because the recombination of *Gdf5-Cre* that occurs in joint primordia during embryonic development is very efficient and specific to cells in the joint^10^. Sagittal sections of the knee at 12 weeks of age and 24 weeks of age without surgical intervention did not show overt abnormalities, suggesting that Piezo channels are not essential for normal articular cartilage development and maintenance (Fig. 1B).

**Fig. 1.**
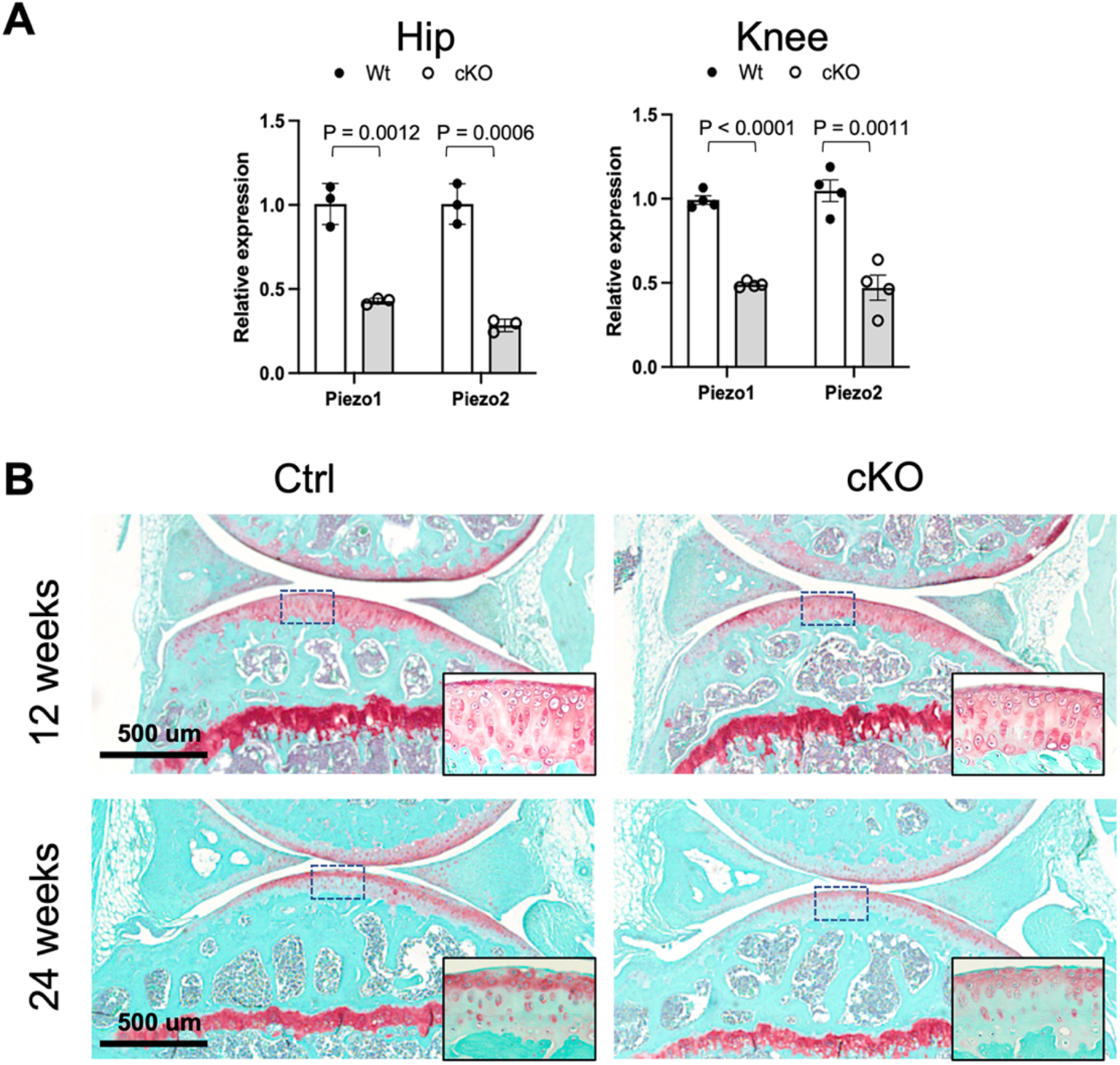
Knee joint morphology of *Piezo1* and *Piezo2* conditional knockout mice. (A) Relative expression of *Piezo1* and *Piezo2* in primary chondrocytes from knee and hip joints of *Piezo1* and *Piezo2* doubly conditional knockout (cKO, *Gdf5-Cre:Piezo1* ^*fl/fl*^*:Piezo2*^*fl/fl*^) mice. assessed. N= 3-4. Two-tailed *t*-test. (B) Safranin-O stained sections of the knee of mice with indicated genotypes and ages. Insets are magnified views of boxed areas. No overt abnormalities are found in cKO mice.

To test whether genetic ablation of Piezo channels affects OA initiation and progression, we surgically induced OA in cKO mice. OA was evaluated 12 weeks after DMM surgery in the tibial plateau according to the OARSI scoring system ^12^. Both cohorts developed moderate to severe OA, but a few cKO mice showed milder OA (Fig. 2A). Mice were also assessed for pain by the incapacitance meter test. We did not find significant differences between the control and cKO groups, suggesting that Piezo channel ablation in the joint does not appear to have beneficial effects on OA-pain (Fig. 2 B).

**Fig. 2.**
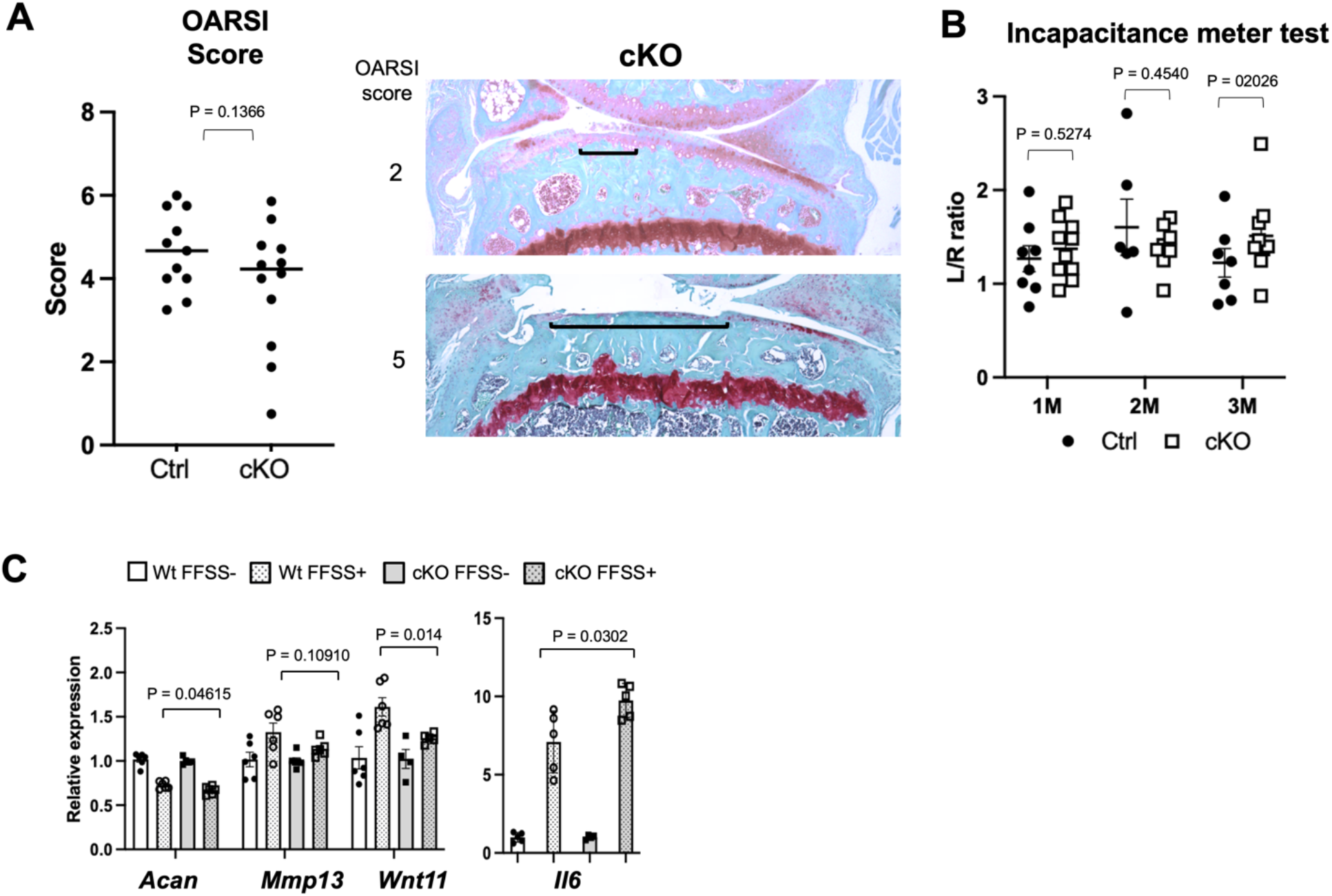
Effects of *Piezo1* and P*iezo2* ablation on OA progression. (A) OA was evaluated on the tibial plateau of *Piezo1* and *Piezo2* doubly conditional knockout (cKO, *Gdf5-Cre:Piezo1* ^*fl/fl*^*:Piezo2*^*fl/fl*^) mice. *Left*. OARSI OA score. Both control and cKO mice develop moderate to severe OA 12 weeks after DMM. The cKO group contains a few samples with mild OA legions. N = 11 for control (Ctrl) and 12 for cKO. Two-tailed Mann-Whitney *U* test. *Right*. Representative images of cKO knees with indicated OARSI scores. Brackets indicate the legion that reaches the calcified cartilage. (B) Pain assessment by the incapacitance meter test. The left-right asymmetry of hindlimb weight bearing (L/R ratio) was quantified. N = 6-10. *t-*test. (C) Gene expression changes in primary articular chondrocytes after 4 hrs of FFSS stimulation. FFSS altered gene expression similarly in both Ctrl and cKO cells. N=5-6, *t-test*.

To investigate the effect of Piezo ablation on the molecular response to mechanical stress, articular chondrocytes isolated from the hip joints of cKO and control mice were exposed to FFSS. The induction of several OA-related genes was assessed. FFSS reduced the expression of *Acan* and increased the expression of *Mmp13, Wnt11*, and *Il6* in both cKO and control cells (Fig. 2 C), although the extent of gene expression change in cKO cells was slightly greater in *Il6* and smaller in *Wnt11* than in control cells.

## Discussion

The critically important roles of Piezo mechanosensing channels in bone formation and homeostasis have been demonstrated ^14^. However, except for a few studies suggesting possible chondroprotective effects of Piezo1 inhibition ^6, 15^, the roles of Piezo channels in cartilage and OA have not been directly demonstrated *in vivo*. Since Piezo channels are activated by strains at injurious levels ^5^ and Piezo1 expression is regulated by inflammatory cytokine signaling ^9^, it is an attractive hypothesis that Piezo channels mediate injurious mechanical stress and thereby promote OA progression. In this study, we attempted rigorously test this hypothesis *in vivo* using a genetic model. Our data show that genetic ablation of *Piezo1* and *Piezo2* has limited effects on cartilage development or OA progression.

Chondrocytes sense mechanical stress through multiple systems. During OA progression, molecules produced by cells in the joint in response to excessive and improper mechanical stimulation are considered to promote the progression of OA. Thus far, our *in vitro* data do not strongly support the role of Piezo channels mediating mechanical stress to the expression of OA-promoting molecules but rather suggest that different mechanosensing mechanisms other than Piezo channels dominantly mediate mechanical stress-induced gene regulation in chondrocytes. However, based on the observation that the Piezo cKO cohort contained samples with milder OA, it is still possible that Piezo inhibition might delay OA progression, if not robustly inhibit it.

## Acknowledgements

We thank the Center for Skeletal Research (P30AR075042) for assistance in histological analysis. This study was supported by the grant, W81XWH1910186, from the U.S. Army Medical Research and Development Command (T.K.).

## Notes

### Competing Interest Statement

The authors have declared no competing interest.

## References

1. Zhao Z, Li Y, Wang M, Zhao S, Zhao Z, Fang J: Mechanotransduction pathways in the regulation of cartilage chondrocyte homoeostasis. J Cell Mol Med 2020; 24:5408–19.

2. Gao W, Hasan H, Anderson DE, Lee W: The Role of Mechanically-Activated Ion Channels Piezo1, Piezo2, and TRPV4 in Chondrocyte Mechanotransduction and Mechano-Therapeutics for Osteoarthritis. Front Cell Dev Biol 2022; 10:885224.

3. McNulty AL, Leddy HA, Liedtke W, Guilak F: TRPV4 as a therapeutic target for joint diseases. Naunyn Schmiedebergs Arch Pharmacol 2015; 388:437–50.

4. O’Conor CJ, Ramalingam S, Zelenski NA, Benefield HC, Rigo I, Little D, et al.: Cartilage-Specific Knockout of the Mechanosensory Ion Channel TRPV4 Decreases Age-Related Osteoarthritis. Sci Rep 2016; 6:29053.

5. Qin L, He T, Chen S, Yang D, Yi W, Cao H, et al.: Roles of mechanosensitive channel Piezo1/2 proteins in skeleton and other tissues. Bone Res 2021; 9:44.

6. Lee W, Leddy HA, Chen Y, Lee SH, Zelenski NA, McNulty AL, et al.: Synergy between Piezo1 and Piezo2 channels confers high-strain mechanosensitivity to articular cartilage. Proc Natl Acad Sci U S A 2014; 111:E5114–22.

7. Mehana EE, Khafaga AF, El-Blehi SS: The role of matrix metalloproteinases in osteoarthritis pathogenesis: An updated review. Life Sci 2019; 234:116786.

8. Molnar V, Matisic V, Kodvanj I, Bjelica R, Jelec Z, Hudetz D, et al.: Cytokines and Chemokines Involved in Osteoarthritis Pathogenesis. Int J Mol Sci 2021; 22.

9. Lee W, Nims RJ, Savadipour A, Zhang Q, Leddy HA, Liu F, et al.: Inflammatory signaling sensitizes Piezo1 mechanotransduction in articular chondrocytes as a pathogenic feed-forward mechanism in osteoarthritis. Proc Natl Acad Sci U S A 2021; 118.

10. Rountree RB, Schoor M, Chen H, Marks ME, Harley V, Mishina Y, et al.: BMP receptor signaling is required for postnatal maintenance of articular cartilage. PLoS Biol 2004; 2:e355.

11. Glasson SS, Blanchet TJ, Morris EA: The surgical destabilization of the medial meniscus (DMM) model of osteoarthritis in the 129/SvEv mouse. Osteoarthritis Cartilage 2007; 15:1061–9.

12. Glasson SS, Chambers MG, Van Den Berg WB, Little CB: The OARSI histopathology initiative - recommendations for histological assessments of osteoarthritis in the mouse. Osteoarthritis Cartilage 2010; 18 Suppl 3:S17–23.

13. Ogawa H, Kozhemyakina E, Hung HH, Grodzinsky AJ, Lassar AB: Mechanical motion promotes expression of Prg4 in articular cartilage via multiple CREB-dependent, fluid flow shear stress-induced signaling pathways. Genes Dev 2014; 28:127–39.

14. Nie X, Chung MK: Piezo channels for skeletal development and homeostasis: Insights from mouse genetic models. Differentiation 2022; 126:10–5.

15. Jones RC, Lawrence KM, Higgins SM, Richardson SM, Townsend PA: Urocortin-1 Is Chondroprotective in Response to Acute Cartilage Injury via Modulation of Piezo1. Int J Mol Sci 2022; 23.

